# Molecular Dynamic Studies on the Interaction of a TatA Oligomer with Tat Translocon Substrates

**DOI:** 10.1101/2025.09.29.679310

**Authors:** Vinh Tran, Terry M. Bricker, Steven M. Theg

## Abstract

The Tat Translocon directly utilizes the Proton Motive Force to transport folded proteins from the n-side to the p-side of energized membranes, targeting the thylakoid lumen of chloroplasts and the periplasmic space of Bacteria and Archaea. In most organisms the Translocon consists of three subunits, TatA, TatB and TatC exhibiting a stoichiometry of ∼20-50/1/1. While TatB/TatC recognize the canonical twin-arginine motif**-**containing signal sequence of substrate proteins, TatA has been hypothesized to interact with TatB/TatC and translocon substrates facilitating their transport across the membrane. TatA from *E.coli* contains a short transmembrane helix near the N-terminus, a longer amphipathic helix and a relatively large unstructured C-terminal domain. While the transmembrane and amphipathic helixes are required for Translocon activity, the C-terminal domain is, in large measure, dispensable. TatA has been hypothesized to form higher-order oligomers in the biological membranes. In this communication we have used 1000 ns-long course-grained molecular dynamic simulations to examine the interactions between a membrane-associated *E. coli* TatA nonamer, alone, and in association with two Tat Translocon substrate proteins, either OEE17 or TorA. In all simulations, either in the presence or absence of substrate, the TatA nonamer markedly thinned the lipid bilayer which may facilitate substrate translocation. The pore of the nonamer was occupied by a phospholipid layer consisting of ∼6 phospholipids in the absence of substrates and ∼11 phospholipids in their presence. Structurally, the amphipathic helix of TatA were observed to exhibit significant conformational flexibility which appears to facilitate TatA-substrate interactions. In the absence of substrate the TatA nonamer was unstable with its radial architecture collapsing in 200-300 ns. In the presence of substrate, however, the radial geometry of the nonamer persists for at least 1000 ns. Interestingly, in the presence of the smaller substrate OEE17, fewer TatA monomers are retained in a radial geometry then observed in the presence of the larger substrate TorA indicating that the molecularity of the TatA oligomer can adjust to the size of the substrate. Specific hydrophilic residues of the TatA amphipathic helixes were found to interact with both substrate molecules, and these form quite stable charge-pair or hydrogen-bonding interactions. While the substrate proteins were initially placed adjacent to the amphipathic helixes of the nonamer, during the simulation trajectories the substrates moved to a more central position adjacent to, and partially entering, the oligomer pore. Concomitantly, the oligomer was observed to lose phospholipids. These latter observations may constitute a glimpse of the initial stages of protein translocation.

## Introduction

The Tat Translocon is an ancient protein transport pathway which is present in all domains of life; Archaea, Bacteria, and Eukarya^1–3^. In Archaea and Bacteria the translocon is found in the cell membrane (and, possibly, the thylakoid membrane in cyanobacteria). In Eukarya the translocon is found uniformly in the plant chloroplast thylakoid membrane, and the mitochondrial inner membrane of plants and some protists. The translocon specifically transports folded proteins across biological membranes ^4, 5^, with the energy for transport being provided directly by the Proton Motive Force (PMF) ^6–8^. An in-depth discussion of *E. coli* Tat Translocon substrates can be found in ^9^. In most organisms the translocon contains three protein subunits, TatA, TatB and TatC, however in some organisms, like *Bacillus*, and most other gram positive bacteria, only a single TatA/TatB component associates with TatC ^10^. In *E. coli* the canonical TatA, TatB and TatC subunits are present with an additional minor TatA-like component, TatE, being expressed at low levels ^11^. The stoichiometry of TatA, TatB and TatC, appears to be ∼25- 50:1:1 ^11^. The large excess of TatA may be related to its hypothesized role in facilitating the passage of substrates across the biological membrane ^12, 13^. A minimal TatABC (1:1:1) subcomplex recognizes and binds substrate proteins which contains a signal sequence bearing a twin-arginine motif proximal to the N-terminus ^14^. Upon substrate binding to the subcomplex, and in the presence of sufficient PMF, additional TatA subunits are recruited, and the signal sequence is processed with the mature substrate protein being translocated across the membrane by an unknown mechanism ^15^. It should be noted that, at least in *E. coli*, TatA can independently associate with substrate proteins in a TatB/C independent process ^16^. NMR structures are available for TatA monomers (PDB: 2LZR ^17^; PDB:2L16 ^18^) and oligomers (PDB:2LZS, ^17^) and for TatB (PDB:2MI2 ^19^), and an X-ray crystal structure is available for TatC (PDB:4B4A ^20^) from *Aquifex aeolicus*.

In *E. coli* TatA is small (89 amino acid residues, 8.4 kDa) and contains a short (fifteen residue) transmembrane helix near the N-terminus, a longer amphipathic helix containing twenty-seven residues and a longer C-terminal random coil domain with forty-two residues (Fig. 1 A-B). The short transmembrane helix is of insufficient length (∼21 Å) to cross the *E. coli* inner membrane (∼40-50 Å, ^21^). This leads to a thinning of the membrane due to a hydrophobic mismatch ^22–24^ which may contribute mechanistically to the transport of substrate proteins across the membrane. The interaction of the amphipathic helix with substrate has been hypothesized to result in additional thinning of the lipid bilayer ^25^ with concomitant membrane destabilization.

**Figure 1.**
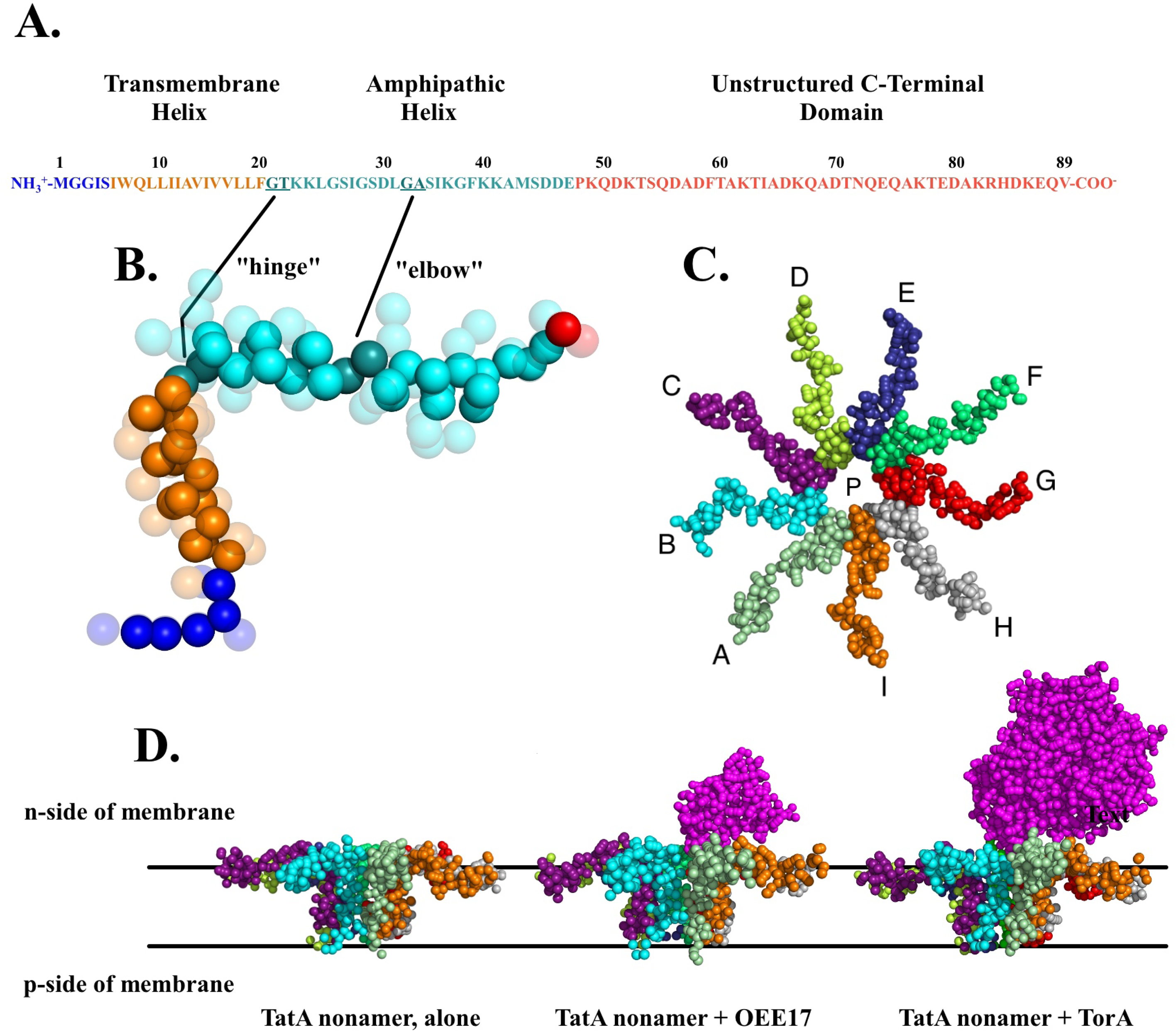
Summary of the systems examined in this communication. A. Amino acid sequence of the 89 amino acid *E. coli* TatA protein. The unstructured N-terminus (residues M1-S5) are shown in blue, the short 15 amino acid residue transmembrane helix (residues I6-F20) is shown in orange, and the amphipathic helix (residues G21-E47) is shown in cyan. Two domains of the amphipathic helix, the “hinge” domain (residues G21 and T22) and the “elbow” domain (residues G33 and A34) are shown in deep cyan and are underlined. Finally, the long, unstructured C-terminus (residues P48-V89) is shown in red. B. Coarse-grained representation of the TatA protein monomer (PDB 2LZR, ^17^). The backbone beads are shown as solid spheres in this representation to highlight the ! helical structure of the transmembrane and amphipathic helixes. The side chain beads are shown as transparent spheres. Color coding is as shown for Fig. 1A. It should be noted that this structure is of a genetically truncated mutant protein ^17^. It had been shown that deletion of the unstructured C-terminal domain had only modest effects on protein transport ^60^. The black lines indicate the locations of the “hinge” and “elbow” domains in correlation to Fig. 1A. C. View of the TatA nonamer (PDB 2LZS, ^17^) as viewed from the n-side of the membrane, which faces the cytoplasm. This representation includes both the backbone and side-chain beads shown as solid spheres. The nine TatA monomers are color coded and labeled A-I. The pore of the nonamer is labeled P and is occupied with membrane lipids (not shown, see Fig. 2). D. View from within the plane of the membrane of the TatA nonamer, alone, the TatA nonamer in association with OEE17 and the TatA nonamer in association with TorA. These images are representative of the different complexes at 10 ns in their trajectories. The solid black lines indicate approximate locations of the phospholipid head groups in the simulated membrane. The n-side (facing the cytoplasm) and the p-side (facing the periplasm) of the membrane are indicated. The membrane lipids, water and Na^+^ and Cl^-^ ions are not shown for clarity.

Cysteine-scanning mutagenesis ^26^ and protein cross linking studies ^27^ have been used to identify the residues in the amphipathic helix which are required for substrate protein translocation. The C-terminal domain is not strictly required for substrate protein transport as it can be removed with only modest effects on Tat translocon function ^26, 28^. The TatA monomers (Fig 1B) appear to form larger oligomeric structures (Fig. 1C). Single particle analysis ^29, 30^, EPR ^31^, and NMR ^17^ structural studies have indicated that TatA can form higher-order radial structures including hexamers, nonamers and dodecamers ^17^.

Molecular dynamic (MD) simulations can provide important insights into the functioning of protein ensembles ^32^. In atomistic MD simulations the motions of each individual atom (including hydrogens) within a molecular system are evaluated. This level of fidelity allows the capture of small, nuanced, interactions. These simulations are particularly valuable when studying processes that depend on atomic precision. However, this resolution comes with a significant computational cost. The number of particles may be very large, and interactions must be calculated for every pair of atoms in a simulated system. As a result, all atom simulations are typically limited to shorter timescales and smaller systems to allow for more a reasonable duration of computation. In our instance, coarse-grain MD simulations were preferred due to the relatively large size of the simulations, their complexity, and limitations due to computing power. In coarse-grained MD simulations adjacent heavy atoms are clustered into single entities termed “pseudo-atoms” or “beads”. This simplifies the evaluation of atomic-scale interactions into broader molecular representations, enabling long timescale simulations of protein dynamics of large ensembles in complex molecular environments which require relatively modest computational resources ^33^. This approach is particularly useful for studying membrane-associated proteins where lipid, water and subunit oligomerization dynamics play key roles in function. By reducing the computational cost of traditional molecular dynamics, coarse-grained MD allows researchers to model large-scale structural rearrangements of the membrane proteins and their lipid environments ^34^. Previous coarse-grained MD simulations suggest that TatA monomers have the potential to associate dynamically to form size-adaptable oligomers, hypothesized to respond to the dimensions of the substrate proteins ^17^. MD simulations have also helped elucidate the role of membrane thinning in facilitating TatA-mediated translocation, highlighting how lipid dynamics influence the translocon’s function ^17, 35^. Finally, MD studies examining the interaction of TatA with the small substrate HiPIP (High Potential Iron-Sulphur Protein) indicate that the mature domain (and signal sequence) of the substrate may interact directly with TatA even in the absence of TatB/TatC. The principal sites of interaction of the mature protein were with the C-terminal half of the TatA amphipathic helix. Unfortunately, the interacting residues were not identified ^16^.

In this communication we have used 1000 ns-long course-grained MD simulations to examine the interactions of a hypothesized membrane-associated TatA nonamer ^17^ in the presence and absence of two Tat substrate proteins, OEE17 (mass = 16.5 kDa from *Pisum satvium*), which is found in the plant thylakoid lumen and which is an extrinsic subunit of Photosystem II ^36, 37^ and TorA (mass = 90.2 kDa from *E. coli*), which is found in the *E. coli* periplasmic space functioning as a trimethylamine-N-oxide reductase ^38^ (Fig. 1D). In our simulations the TatA nonamer, both alone and in the presence of substrates significantly thin the biological membrane as suggested by other investigators ^17, 22, 24, 29, 35^. Additionally, both in the presence and absence of substrates, the amphipathic helixes exhibit significant conformational flexibility. In the presence of substrates this flexibility facilitates the interaction of substrate residues with specific polar residues of the amphipathic helixes. Our findings indicate that the TatA nonamer is quite unstable in the absence of substrate with its radial geometry (^1^Fig. 1C) usually collapsing in < 300 ns. In the presence of substrate, the radial geometry of the oligomer persists for at least 1000 ns. Interestingly, the number of TatA monomers which are retained in the radial geometry while in association with the substrate varies depending on the size of the transported proteins. In the presence of OEE17, six of the TatA monomers are usually retained in the radial architecture while, when associated with TorA eight monomers are usually retained. Additionally, specific residues of the TatA amphipathic helixes interact with both substrate molecules, and these form apparently stable (100s of ns timescale) charge-pair and/or hydrogen-bonding interactions. The interacting TatA residues we identify are highly congruent with residues identified earlier by cysteine-scanning mutagenesis in *E. coli* to facilitate Tat Translocon activity ^17^, with chemical crosslinking studies which identified residues interacting with the Translocon substrates SufI and TorA ^39^, and experiments in higher plants which identified Tha4 (higher plant homologue of TatA) residues which interact with OEE17 ^27^. These authors hypothesized that these interactions may facilitate substrate translocation. Our MD simulations support this hypothesis. The nonamer pore was occupied by a layer of phospholipids as reported earlier ^17^ consisting of ∼6 phospholipids in the absence of substrate and ∼11 in the presence of either OEE17 or TorA. In Rodriguez et al. ^17^ the lipid layer was described as being “distorted”. Interestingly, in our simulations, the lipids present in the TatA oligomeric pore appeared to form a lipid monolayer. Finally, In our simulations we initially placed the substrates adjacent to the TatA amphipathic helixes. During the trajectory time-course of the simulations the substrates moved towards and partially penetrate the nonamer pore. Concomitant to the movement of substrate, phospholipids are lost from the pore. These latter observations may constitute the initial stages of protein translocation.

## Materials and Methods

Three initial atomistic models of the *E. coli* TatA nonamer were constructed using PYMOL ^40^. These were, 1. TatA nonamer, alone (PDB: 2LZS, ^17^), 2. the TatA nonamer associated with OEE17 (PDB:1NZE, ^41^), and, 3. the TatA nonamer associated with TorA (PDB: 1TMO, ^42^) (Fig. 1D). These structures were obtained from the RCSB Protein Data Bank (www.rcsb.org) and imported into PYMOL. The OEE17 and TorA substrates, individually, were positioned proximal (5-10 Å) to the TatA subunits G and H (Fig. 1C), and adjacent to the pore region of the nonamer. The positioning of the substrate molecules was offset with respect to the radial geometry of the TatA nonamer (Fig. 1D). These three configurations were exported as pdb files and input, individually into the CHARMM-GUI web-based interface ^43^ and each was used to construct a “course-grained” model of the membrane system. The “Martini Bilayer Maker” option was used in conjunction with the martini22p forcefield ^44^. This forcefield incorporates both polarized water and charged amino acid residues. Each model configuration was uploaded individually and conditions were kept consistent between the three configurations. All simulations were conducted at pH 7.0 and at a 150 mM NaCl concentration. Initially, the protein components were oriented by adjusting the Z axis with respect to the membrane allowing lipids to populate the nonamer pore. The systems measured 200 x 200 Å in the x- and y axis.

Due to large differences in substrate size, the z-axis was adjusted to 100 Å in the absence of substrate, 150 Å in the presence of OEE17, and 250 Å in the presence of TorA. The lipid composition of the modeled *E. coli* inner membrane consisted of 80% POPE (1-palmitoyl-2- oleoyl-sn-glycero-3-phosphotidylethanolamine), 15% POPG (1-palmitoyl-2-oleoyl-sn-glycero- 3-phosphotidylglycerol) and 5% cardiolipin (1,3-bis[1’,2’-dioleoyl-sn-glycero-3-phospho]-sn-glycerol).

The preliminary minimization and equilibration steps were run using the default parameters of CHARMM-GUI ^44^, at a temperature of 303 K and a pressure of 1 bar. The CHARMM-GUI output files were downloaded and the MD simulations were run in GROMACS version 2020.3 (DOI: 10.5281/zenodo.3923644). The “step7_production.mdp” file was modified to run the MD calculations for 50,000,000 steps or the equivalent of 1000 ns (20 fs/step). The GROMACS output files (.gro, .xtc, .tpr, etc.) were visualized and analyzed using PYMOL and VMD ^45^. Five independent simulations, each, were performed for the TatA nonamer, alone, the TatA nonamer associated with the substrate OEE17, and the TatA nonamer associated with the substrate TorA. Sequence alignment was performed using Clustal Omega ^46^. All simulations were performed on a Silicon Mechanics Workform GX-RT2 workstation.

## Results and Discussion

Fig. 2A illustrates a membrane embedded TatA nonamer, alone, at 10 ns in a simulation trajectory. Shown is a transect approximately bisecting the nonamer (note that some foreground TatA monomers and lipids were hidden in this view for clarity). We emphasize that all the of the simulations, in both the absence and presence of substrate, exhibit this same basic architecture at 10 ns. Initially, the short transmembrane helixes of all of the TatA monomers are embedded in the lipid bilayer and their amphipathic helixes largely rest on the ^2^n-side surface of the membrane. The pore (Fig. 1C) of the nonamer is occupied by a phospholipid layer and water as described previously ^17^. The short length of the transmembrane helixes, 15 amino acid residues, leads to a hydrophobic mismatch ^22–24^ which distorts the lipid bilayer, leading to membrane thinning; this effect is amplified by displacement of n-side polar head groups by the amphipathic helixes. Outside of the nonamer, the membrane is 43.5 ± 3.8 Å thick. This simulated value compares quite favorably to the *E. coli* cell membrane bilayer thickness of 40-50 Å as determined experimentally by small-angle X-ray scattering, small-angle neutron scattering ^47, 48^ and earlier MD simulations ^49^ The phospholipid within the nonamer pore forms a layer with a thickness of 21.3 ± 5.1 Å (Fig. 2B). This is similar to that observed in an earlier MD study (see Fig. 7 in ^17^). In our simulations of the TatA nonamer, alone, the nonamer pore contained on average 6.6 ± 1.3 phospholipids while in the presence of substrates (TatA nonamer + OEE17 and TatA nonamer +TorA simulations) an average of 11.5 ± 1.0 phospholipids were ^3^identified. The functional significance, if any, of this observation in unknown. Surprisingly, the phospholipid layer present in the nonamer pore nearly uniformly appears to be organized as a monolayer (Fig. 2). Across all simulations, 95% of the phospholipid polar head groups face the p-side of the membrane. In biological systems, phospholipid monolayers are exceedingly rare, being only observed at phase-interfaces, such as on the surface of lipid droplets and in tear films ^50, 51^. It is unclear if our result is significant, possibly contributing to the translocation mechanism, or is an artifact of the Martini Bilayer Maker-GROMACS simulation. In earlier work the hydrophobic layer occupying the pore was observed to be approximately ½ the thickness of a normal lipid bilayer with the phospholipids being described as “distorted” ^17^. Our observations are consistent with this earlier result. Regardless of the precise nature of the lipid layer occupying the pore (lipid monolayer vs. distorted lipid bilayer), we hypothesize that this phospholipid layer forms a barrier to prevent the free diffusion of water, ions and other small molecules across the membrane while facilitating a more energetically feasible pathway for proteins to cross the cell membrane bilayer.

**Figure 2.**
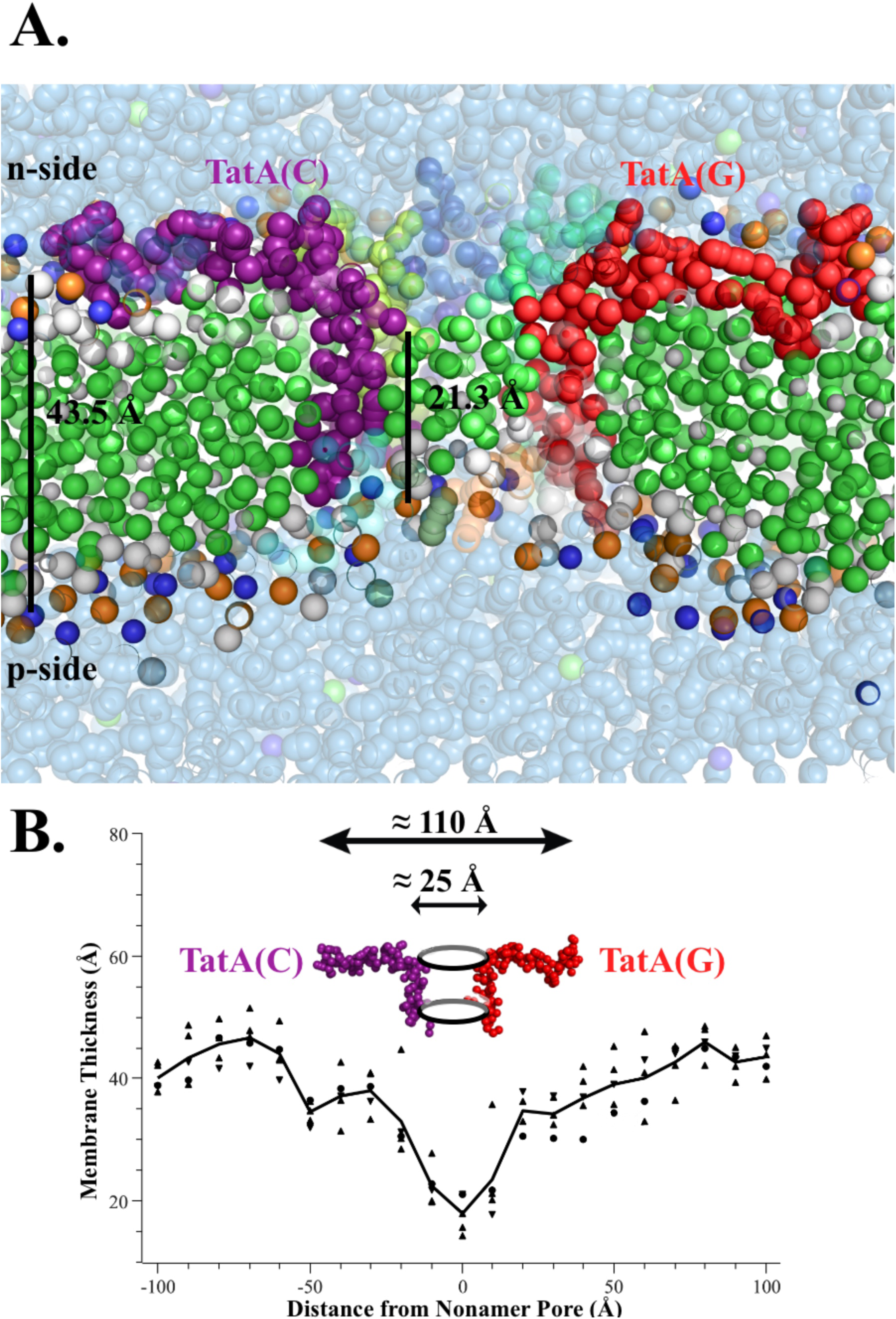
Closeup view of a the TatA nonamer. A., The TatA monomers are color-coded as in Fig. 1C. The membrane phospholipids are shown. The beads representing their aliphatic carbons are shown in green and the beads representing the polar head group atoms are shown in orange (phosphorus), blue (nitrogen) and white (oxygen). TatA(C) and TatA(G) are labeled. Polarized water is shown as transparent pale blue spheres. It should be noted that four of the TatA monomers (TatA(B), TatA(A), TatA(I), and TatA(H)) were hidden in this view. The thickness of the lipid bilayer to the outside of the nonamer is indicated by the long black line (43.5 Å) and thickness of the apparent lipid monolayer within the pore of the nonamer (short black line, 21.3 Å) are indicated. This view is after 10 ns of simulation time and is essentially identical in all fifteen simulations performed either in the absence or presence of substrates. Substrate proteins, and ions are not shown for clarity. B. Membrane thickness as a function of distance from the nonamer pore. A transect was taken through the center of the pore and membrane thickness measurements were taken at 10 Å intervals. Plotted are the results for all five TatA nonamer, alone simulations at 10 ns in the trajectory. Similar patterns were observed in the presence of substrate (TatA nonamer + OEE17 and TatA nonamer+ TorA simulations). The positions of Tat(A) and Tat(G) and location of the pore are shown as are the approximate dimensions of the nonamer and pore.

The amphipathic helix of TatA has been hypothesized to interact with substrate as a prelude to translocation. The helix is quite asymmetric with the hydrophobic face subtending 120-140° and the hydrophilic face subtending 240-220° of the helical wheel (Fig. S1). Examination of the simulations both in the absence and presence of substrates (at 10 ns) indicates that the residues on the hydrophobic face preferentially interact with the aliphatic carbons of the membrane lipids while residues at the boundary between the hydrophobic face and the hydrophilic face (T22, S35, K40, D46 and E47) interact preferentially with the phospholipid head groups on the n-side of the membrane. These amphipathic helixes contain two regions which appear to confer conformational flexibility; the “hinge”domain (G21-T22) ^17^ and the “elbow region” (G33-A34) which could facilitate TatA-substrate interaction (Fig. 1A and 1B). The presence of glycyl residues is known to destabilize amphipathic helices ^52^. This hypothesized conformational flexibility of the amphipathic helixes was explicitly observed in our simulations. Fig. 3 illustrates a simulation trajectory for all of the TatA subunits from a single TatA nonamer, alone. While initially, at 10 ns, the amphipathic helixes largely rest on the n-side surface of the membrane, conformational changes, involving both the “hinge” and “elbow” regions occur throughout the trajectory. Interestingly, three addition glycyl residues are present in the amphipathic helix of TatA, G26, G29 and G37,; only modest additional flexibility was observed at these positions. In Fig. 3, for simplicity, the membrane boundaries are presented as static lines (p-side and n-side). In the actual simulations this was clearly not the case as the lipid bilayer exhibited significant conformational adaptation to the structural mobility of both the amphipathic helixes and the position of the transmembrane helix. It should be emphasized that similar flexibility of the amphipathic helix was uniformly observed in all simulations regardless of the presence or absence of substrate, particularly at shorter simulation times. In the presence of substrates, specific residues of the amphipathic helixes were observed to associate, by either apparent charge-pair or apparent hydrogen-bonding interactions, with substrate residues (see below). Once formed, these tended to be quite stable, limiting further conformational changes of the substrate-associated amphipathic helixes.

**Figure 3.**
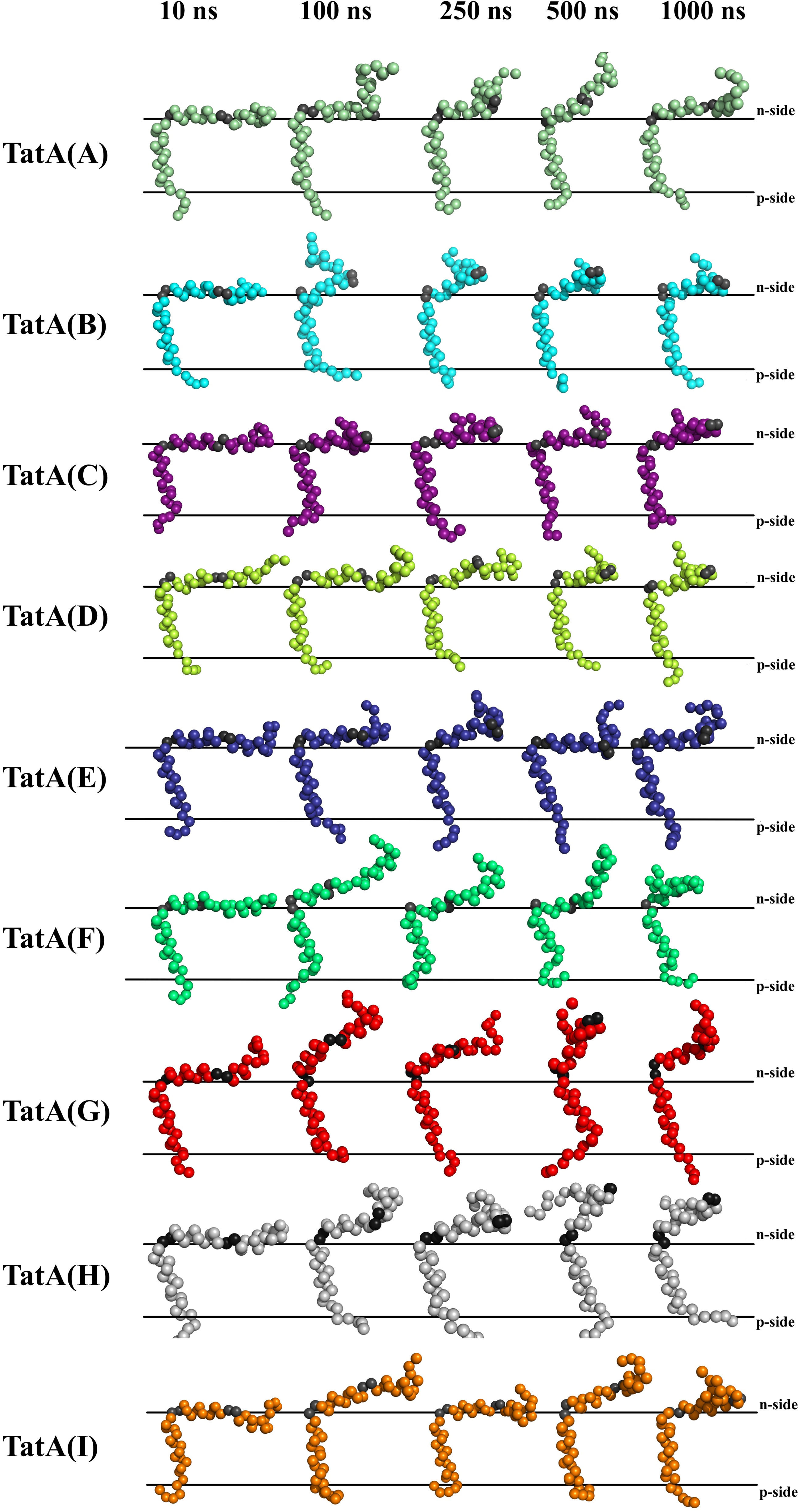
Conformational flexibility of the amphipathic helix during a simulation trajectory of TatA nonamer, alone. Shown are snapshots of the nine TatA monomers within the nonamer taken at 10, 100, 250, 500 and 1000 ns. Color coding for the subunits are as in Fig. 1C. Two domains, the “hinge” (G21-T22, to the left) and the “elbow” (G32-A33, to the right) are shaded dark gray. Only the backbone beads are shown for clarity. The approximate boundaries of the membrane are shown by black lines and the n-side and p- side of the membrane are labeled. Significant flexibility is observed at both the “hinge” and “elbow” regions. The timing and extent of these conformational changes is different for each monomer. It must be emphasized that very similar conformational changes are observed for the TatA monomers in all three of the models simulated. Water, membrane lipids and ions are hidden for clarity.

Fig. 4 and B illustrates the TatA nonamer, alone, the TatA nonamer + OEE17 and the TatA nonamer + TorA at 10 ns and 1000 ns in typical simulations. In Fig. 4A the complexes are viewed from the p-side of the membrane and in Fig. 4B they are viewed from within the plane of the membrane. In both instances, the amphipathic helixes have been hidden for clarity and only the backbone of the transmembrane helixes are shown. Qualitatively, in the majority of simulations in the absence of substrates (TatA nonamer, alone) the radial architecture of the nonamer essentially collapses (Fig. 4A.1 and 4B.1). This occurs in 200-300 ns. This instability in the absence of substrates was noted in an earlier MD study ^17^. However, in the presence of either substrate the radial architecture persists for at least 1000 ns. This observation indicates that the presence of substrates stabilizes the overall structure of the TatA oligomer. Additionally, the size of the substrate influences the molecularity of the TatA oligomer (Fig. 5). The larger substrate, TorA, maintains on average, significantly more TatA monomers in the oligomeric complex at 1000 ns (7.6 ± 0.9, n = 5) monomers than the smaller substrate OEE17 (6.0 ± 1.2, n =5) monomers. Finally, during the time-course of the simulations the substrates move from the periphery of the TatA nonamer (at 10 ns), where they are proximal to the amphipathic helixes (Fig. 1D, Fig. 4A.2 and 4.A.3) to a more centralized location, where they are adjacent to, and in most instances (nine of ten simulations), partially penetrate the oligomer pore (at 1000 ns) (Fig. 4B.2 and 4B.3). In these instances, both the OEE17 and TorA substrates penetrate >5 Å below the phospholipid head groups on the n-side of the membrane. Water, which at 10 ns occupies the n-side of the pore (Fig. 2), is partially or completely displaced by this substrate movement. As the substrate moves to partially occupy the nanomer pore, some of the phospholipids which initially (at 10 ns) occupy the pore are displaced to the bulk membrane (Fig. S2). It is tempting to speculate that this movement of substrates toward the oligomer pore, their partial penetration into the pore cavity with the displacement of water and pore phospholipids may represent the first stages of substrate translocation.

**Figure 4.**
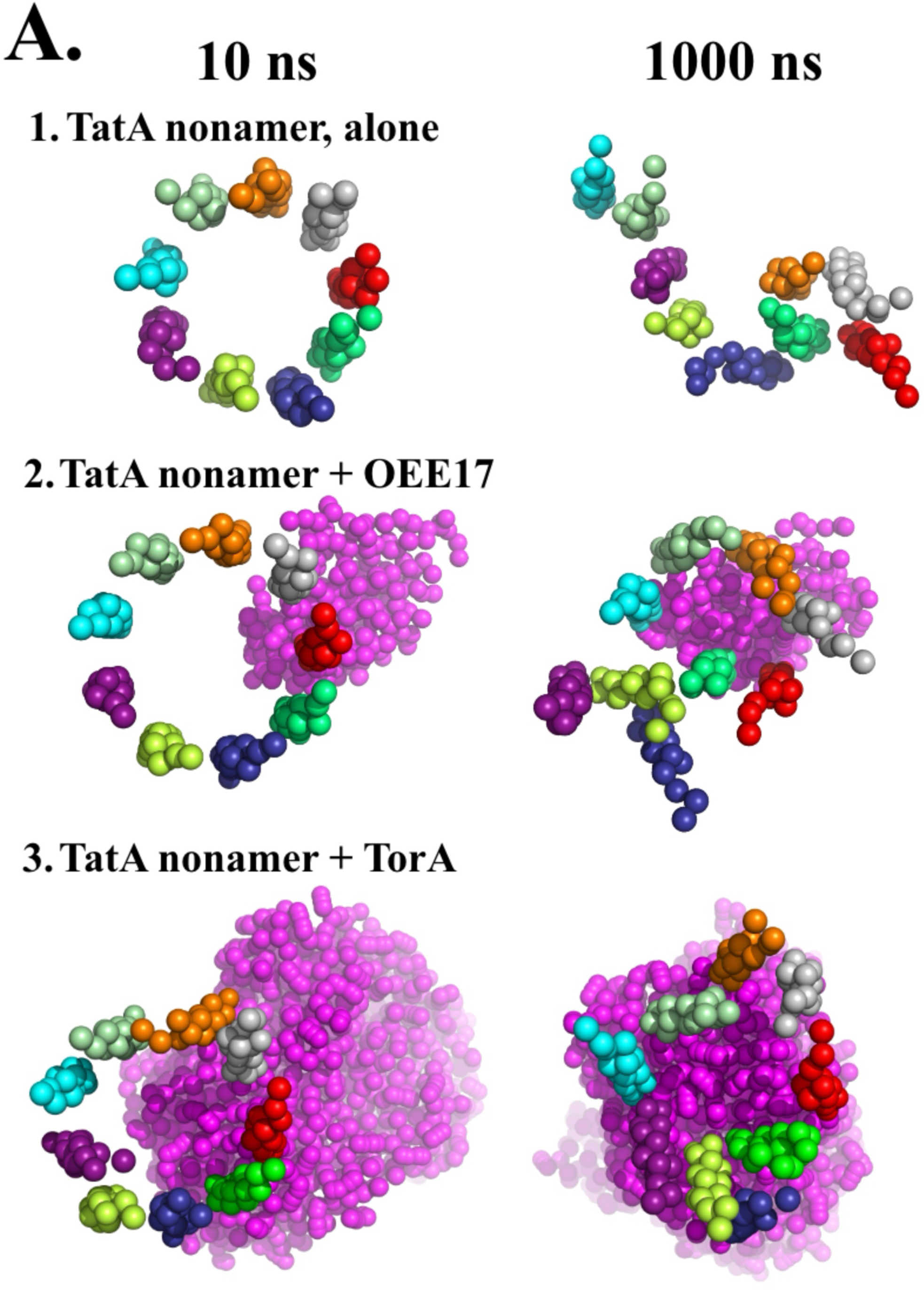
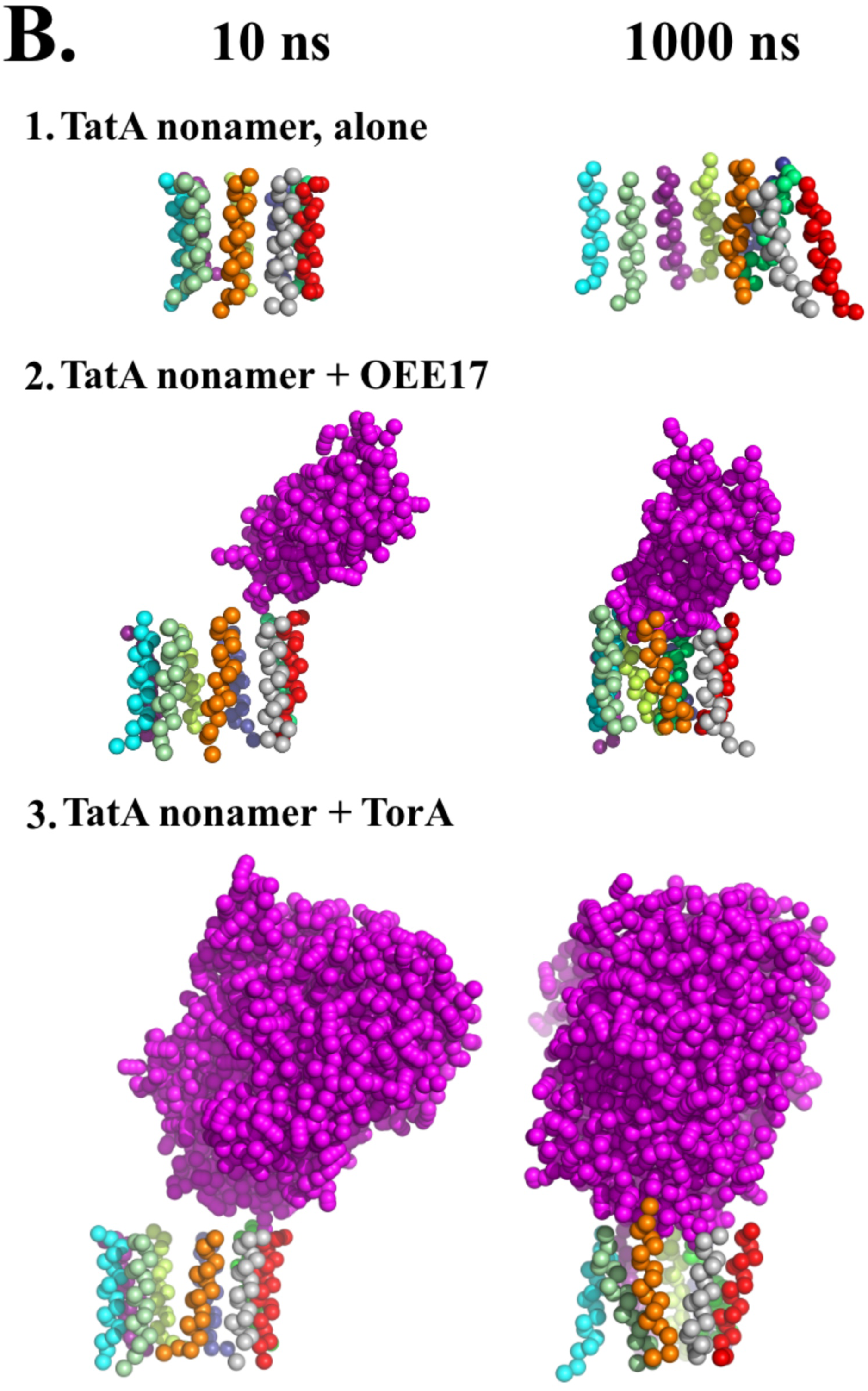
Stability of the radial geometry of the TatA nonamer in the absence or presence of substrates. Shown are the results from typical simulations. Only the backbones of the transmembrane helices are shown (color coded as in Fig. 1C) for each TatA monomer with the amphipathic helixes being hidden for simplicity. Additionally, the water, membrane lipids and ions are hidden in these representations. A. View of the TatA nonamer (and substrates) from the p-side of the membrane. Substrate proteins, which are located on the n-side of the membrane, are shown in magenta. The configuration of these components after 10 ns of simulation are shown to the left and are essentially identical for each of the five simulations performed on each model (Fig. 1D) except for the presence or absence of substrates. To the right is shown the configuration of these components after 1000 ns. In Fig. 4.A.2, the three TatA monomers which were largely excluded from the radial geometry after 1000 ns were TatA(C), TatA(D) and TatA(E). In Fig 4.A.3 the single TatA monomer excluded from the radial geometry was TatA(E). In Fig.4.A.2 and 4.A.3 movement of the substrate to a more central position within the nonamer is evident. B. View of the TatA nonamer (and substrates) from within the plane of the membrane both at 10 ns (left) and 1000 ns (right). In Fig.4.B.2 and 4.B.3 movement of the substrate to a more central position within the nonamer and partial penetration of the substrate into the nonamer pore is evident.

**Figure 5.**
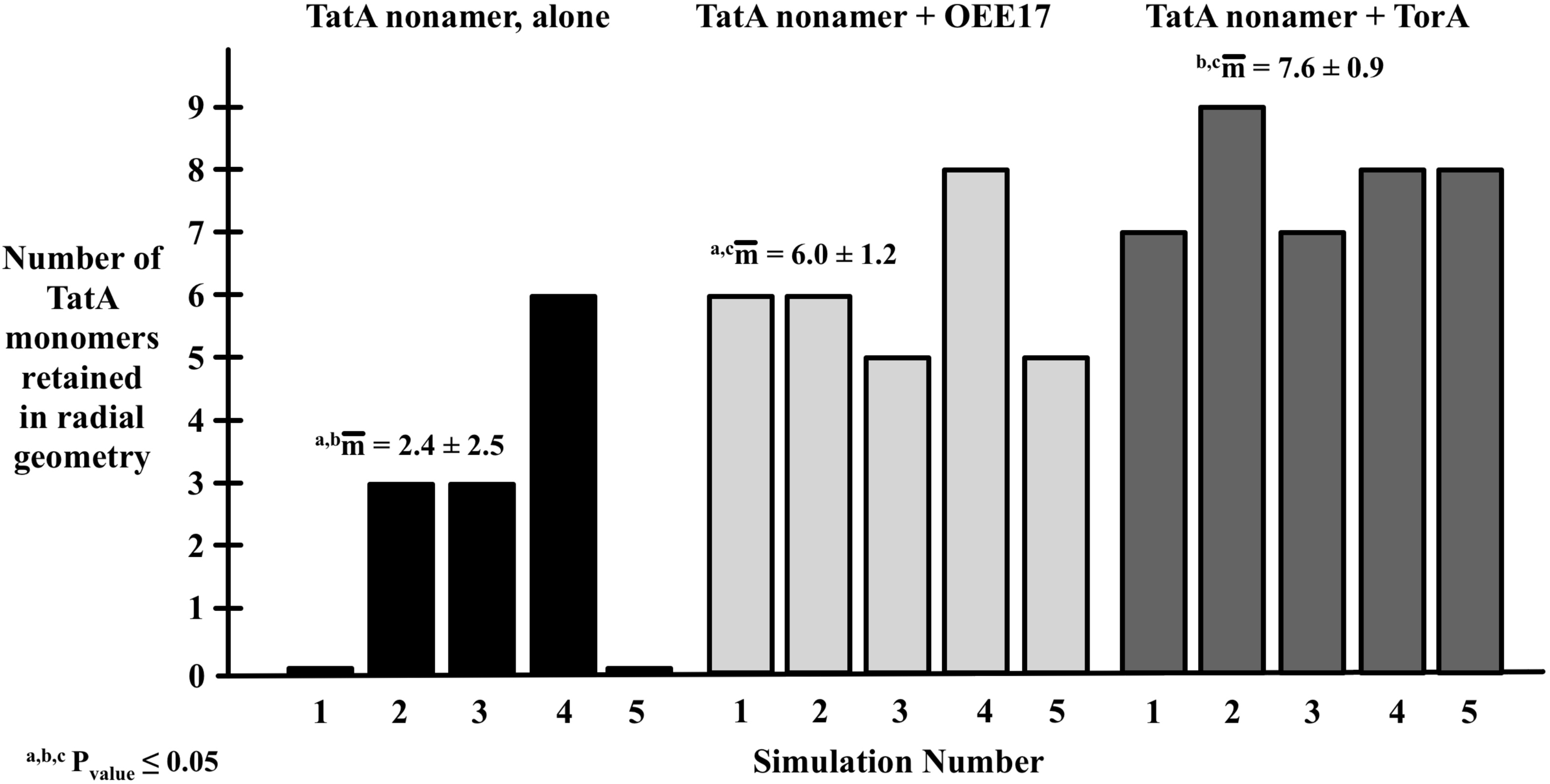
Summary of the retention of the radial geometry of the TatA nonamer in the presence and absence of substrate for all simulations. The number TatA monomers retained in the radial geometry is shown to the right, while the simulation number for each model (as shown in Fig. 1D) is indicated below. The mean values and standard deviations for each model type are shown. Two-way t-tests were performed for each pair of model types. All models exhibited significantly different numbers of TatA monomers within the observed radial geometries, and these were correlated to presence vs. absence of substrate and to the size of the substrate molecules.

During the course of the TatA nonamer + OEE17 and TatA nonamer + TorA simulations, specific residues of the TatA amphipathic helixes were observed to interact with the substrate residues by ^4^apparent hydrogen-bond and apparent charge-pair interactions. These are identified in Table 1. Over the ten substrate-containing simulations performed, seventy-one interactions were identified at 1000 ns; seventeen were apparent hydrogen-bonding ineractions and fifty-four were apparent charge-pair interactions. These formed at various times (10s to 100s of ns) during the the course of the simulations, and the majority (87%) of these were stable over the last 200 ns of the simulations. The distances between the interacting residues were measured during this 200 ns window to determine the average bond length of the interactions (Table 1). The bond length of the apparent charge-pair interactions was 2.6 ± 1.8 Å which is consistent with these being moderate-strong interactions ^53^, while the bond length of the apparent hydrogen-bond interactions was 4.1 ± 2.0 Å which is consistent to these being weak-moderate interactions ^54, 55^.

**Table 1.**
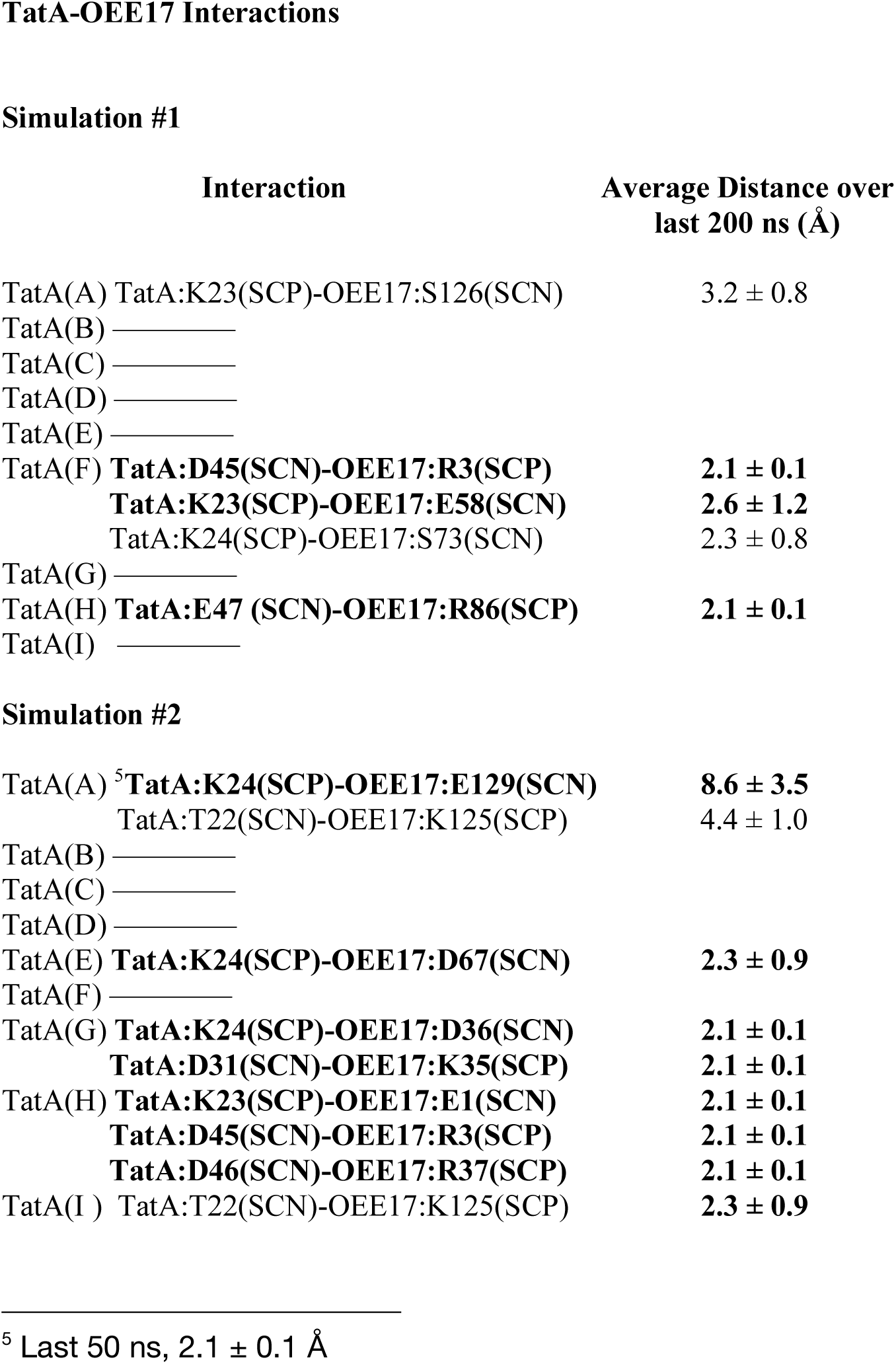
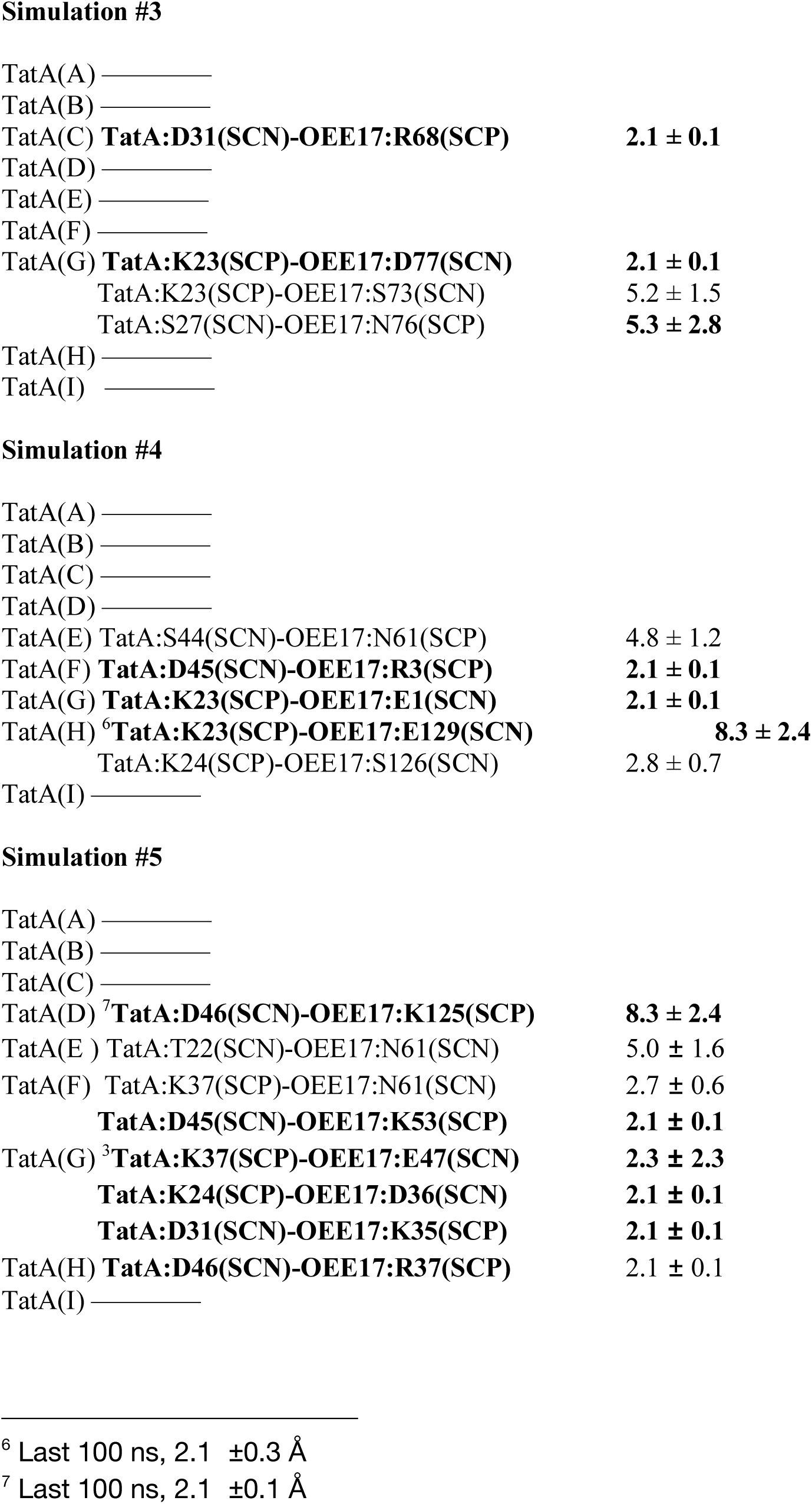
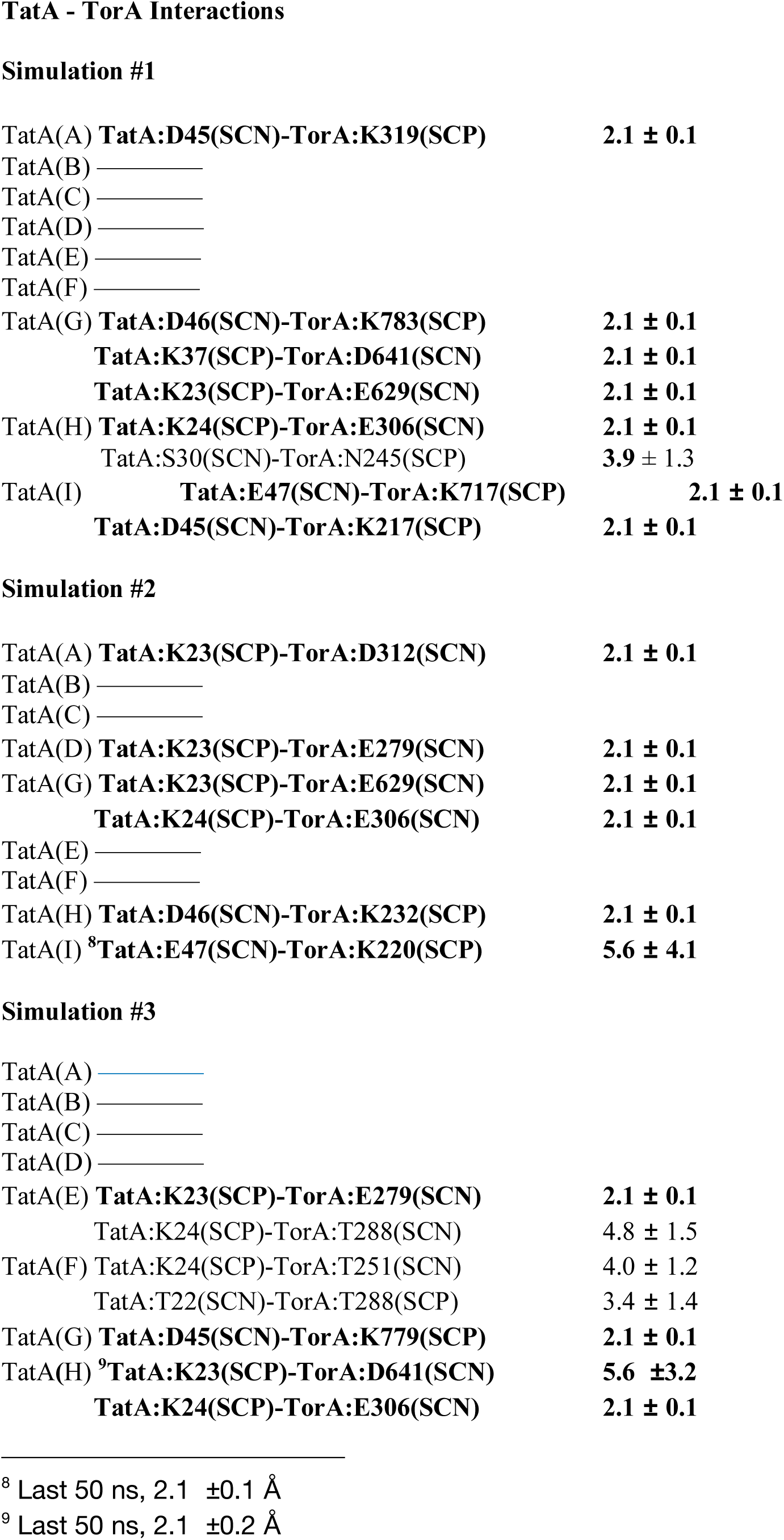
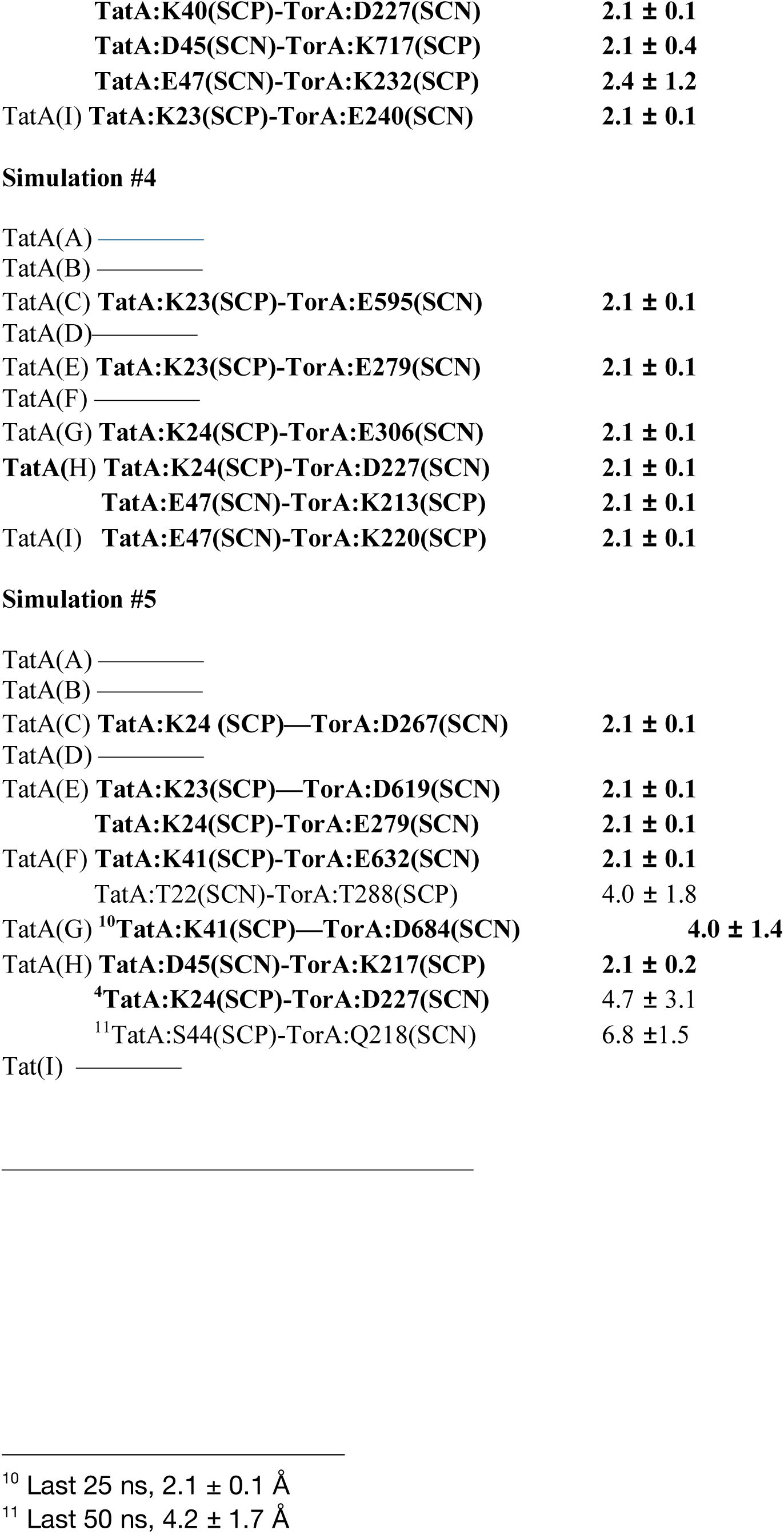
Observed TatA-Substrate Interactions at 1000 ns. Individual simulations were examined and apparent interacting residues (≤ 3.5 Å) were determined by inspection. Within a simulation, individual TatA monomers are identified by their code ((Fig. 1C (A-I)) in parentheses. The interacting residues are identified by the single letter amino acid code. The individual interacting “placeholder” beads (SCP and SCN), as identified in the PDB output file from “Martini Membrane Maker” (files “step5_charmm2gmx.pdb”), are shown. Apparent hydrogen-bond interactions are shown in regular text while apparent charge-pair interactions are shown in bold text. Additionally, the average distance between the interacting residues over the last 200 ns of the simulations are shown

In many instances, large conformational changes were observed in the interacting amphipathic helixes. This is illustrated in Fig. 6A in which the snapshots of the trajectory of the TatA(F):D45-OEE17:R3 interaction (Table 1, TatA nonamer-OEE17 Simulation #1) is shown. At 10 ns these residues are 47.1 Å apart while at 750 ns they are 2.2 Å apart, a distance which remains essentially invariant over the last 250 ns of the simulation. In Fig. 6B, distances within this trajectory are shown in red and two other charge-pair interacting residues TatA(H):E47-OEE17:R86 (blue), with an initial distance between these residues of 59.9 Å, and TatA(F):K23- OEE17:E58 (green), with an initial distance between these two residues of 9.9 Å, are shown.

**Figure 6.**
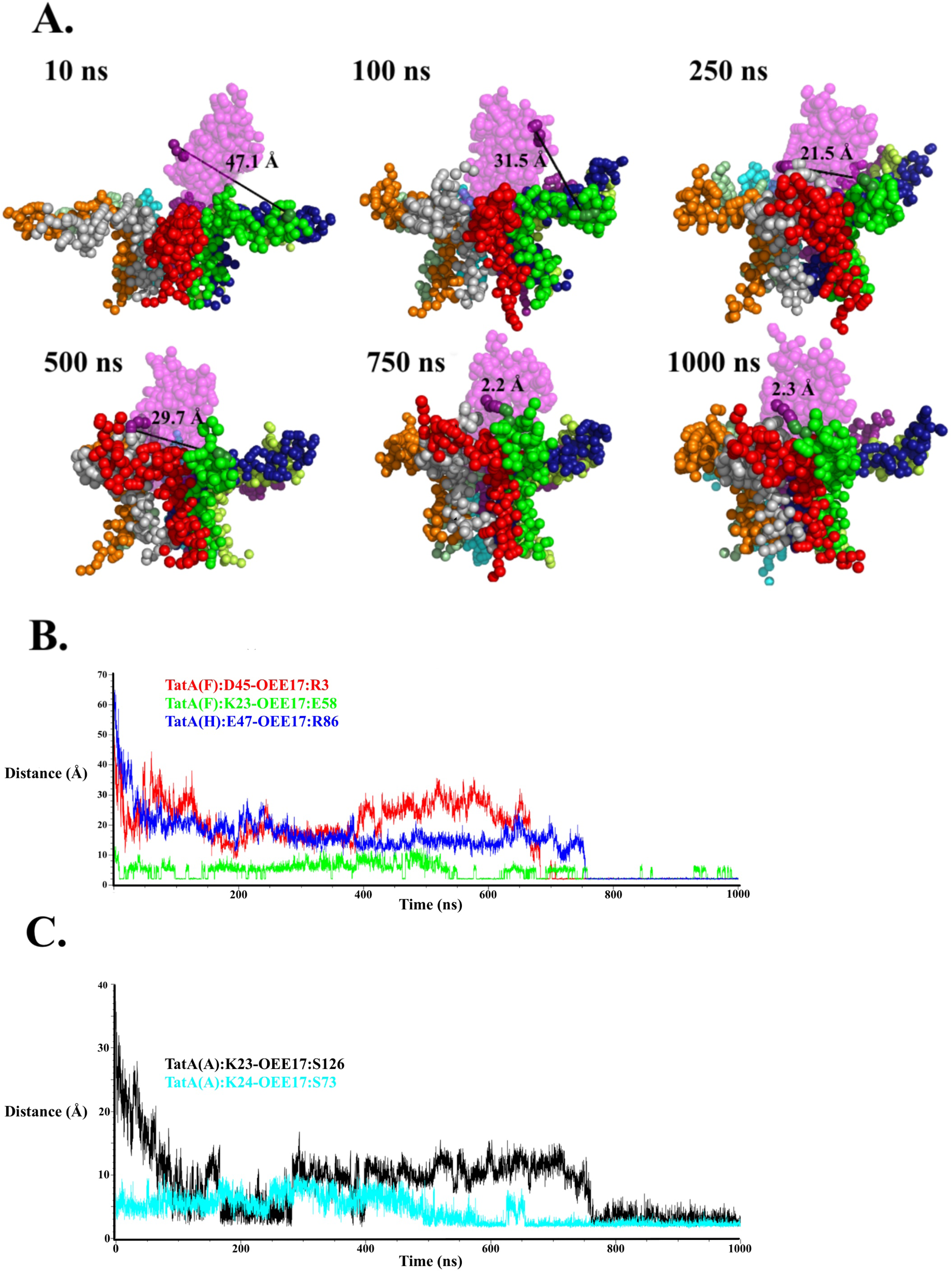
Example of conformational changes observed during the interaction of the TatA nonamer with the substrate OEE17. The results in this figure are from a single TatA nonamer-OEE17 simulation. A. Individual frames from a trajectory highlight the interaction between the residues TatA(F):D45 and OEE17:R3. The residue TatA(F):D45 is shown in dark green while the residue OEE17:R3 is shown as deep magenta. The distance between these two residues is shown in Å and indicated by a black line. Individual TatAs are color-coded as in Fig 1C and represented as solid spheres. The substrate OEE17 is colored magenta and is represented by transparent spheres. B. The trajectory of three different apparent charge-pair interactions are shown in this distance vs. time plot. The interaction of TatA(F)D45-OEE17:R3, which is highlighted in A., above, is shown in red. The interaction of TatA(F):K23-OEE17:E58 is shown in green and that of TatA(H):E47-OEE17:R86 is shown in blue. C. The trajectory of two different apparent hydrogen-bond interactions are shown in this distance vs. time plot. The interaction of TatA(A):K23-OEE17:S126 is shown in black and the interaction of TatA(A):K24-OEE17:S73 is shown in cyan. Note different Y-axis scales in B. and C.

These interactions yield similar results with terminal distances between these residue pairs of about 2 Å. Fig. 6C illustrates similar conformational changes between residues interacting via apparent hydrogen-bonds. The residue pair TatA(A):K23-OEE17:S126 (black trace) are initially separated by 37.4 Å while TatA(A):K24-OEE17:S73 (cyan trace) are separated by 5.8 Å. These exhibited terminal distances during the last 200 ns of the simulation of 3.2 and 2.3 Å, respectively (Table 1, TatA-OEE17 Interactions, Simulation #1). For both of these interactions the range of distance values observed over the last 200 ns is broader than those observed for the residue pairs interacting via apparent charge-pairs (Fig. 6B). This yields a larger standard deviation for these measurements and implies that these interactions are generally weaker than observed for the charge-pairs. All of the OEE17- and TorA-containing simulations exhibit analogous results to those explicitly shown in Fig. 6 and a summary of these interactions are shown in Table 1.

Table 2 summarizes the number of observed interactions between TatA and the substrates OEE17 and TorA with respect to the position of the residues within the TatA amphipathic helix. With respect to total interactions, two regions appear to be particularly important. The residues T22-K24 account for thirty-seven of the observed interactions (52%), while the residues D45- E47 accounts for twenty-one of the observed interactions (30%). Only minor differences were observed when considering each substrate individually. This suggests that substrate size does not significantly influence the pattern of TatA-substrate interactions.

**Table 2.** Summary of the Amphipathic Helix Amino Acid Residues which interact with the Tat Translocon Substrates OEE17 and TorA.

| **TatA Residue** | **OEE17 Interactions** | **TorA Interactions** | **Total Interactions** |
| --- | --- | --- | --- |
| **T22** | 3 | 2 | 5 |
| **K23** | 7 | 10 | 17 |
| **K24** | 6 | 10 | 16 |
| **S27** | 1 | 0 | 1 |
| **S30** | 0 | 1 | 1 |
| **D31** | 3 | 0 | 3 |
| **K37** | 2 | 1 | 3 |
| **K40** | 0 | 1 | 1 |
| **K41** | 0 | 2 | 2 |
| **S44** | 1 | 1 | 2 |
| **D45** | 4 | 5 | 9 |
| **D46** | 4 | 2 | 6 |
| **E47** | 1 | 5 | 6 |
| **Total** | 32 | 40 | 72 |

If the interacting residues that we have observed are important for the interaction of TatA with its substrates, it would be expected that mutagenesis of these residues would compromise Tat Translocon activity. An earlier report indicates that this the case. Greene et al. ^26^ performed cysteine scanning mutagenesis on the amphipathic helix residues of *E. coli* TatA. All of the residues in the region G21-M43 were mutagenized individually to cystinyl residues and Tat Translocon activity was monitored by measuring trimethylamine-N-oxide reductase activity (TorA). In Fig. 7 we have redrawn their results (Greene et al. ^26^, their Fig. 2B) and compared these to the interactions we observed between TatA and the two substrates (Table 2). In large measure, our results correlate with their results ^26^. In the regions where they observed maximal loss of activity due to mutagenesis (G21-L25, G29-G33, and I36-K40) we observed interacting residues (particularly in their domain G21-L25). Mutagenesis of the two glycyl residues at the “hinge” and “elbow” regions (G21 and G29, respectively) also led to a complete loss of TatA translocation activity ^26^ which appears to indicate that the increased mobility of the “hinge” and “elbow” regions are required for Tat translocation and correlates with our observation of increased mobility of the amphipathic helix at these positions (Fig. 3). It should be noted that mutagenesis of seryl (and possibly threonyl residues ^56, 57^) to cystinyl residues in ^26^ would not necessarily be expected to lead to complete loss of activity ^58^. The -SH functional group of cysteine would have similar, although weaker, hydrogen bonding tendencies as the -OH group on serine (and possibly, threonine). This may at least partially explain the relatively high activity of the mutants S27C, S30C, and S35C observed in these studies ^26^. Unfortunately, a number of the TatA residues which we observed to interact with the substrates were not tested in this mutagenesis study including, S44, D45, D46 and E47. Chemical crosslinking studies in *E. coli* ^39^ have identified one of these residues, D46, as interacting with two Tat Translocon substrates SufI and TorA. Additionally, residues in the domain S27-F39 also interacted with these two substrates (Fig. S3). We also observed several of these residues interacting with OEE17 and TorA (specifically S27, S30, D31 and K37). Additional MD studies examining the interaction of TatA with the small substrate HiPIP (High Potential Iron-Sulphur Protein) indicated that the mature domain (and signal sequence) of the substrate may interact directly with TatA even in the absence of TatB/TatC. The principal sites of interaction of the mature protein were with the C-terminal ½ of the TatA amphipathic helix. Unfortunately, the interacting residues were not identified ^16^. Nevertheless, this result is consistent with our observation that residues in the domain D45-E47 which lie in the C-terminal half of the TatA amphipathic helix are involved in TatA-substrate interactions.

**Figure 7.**
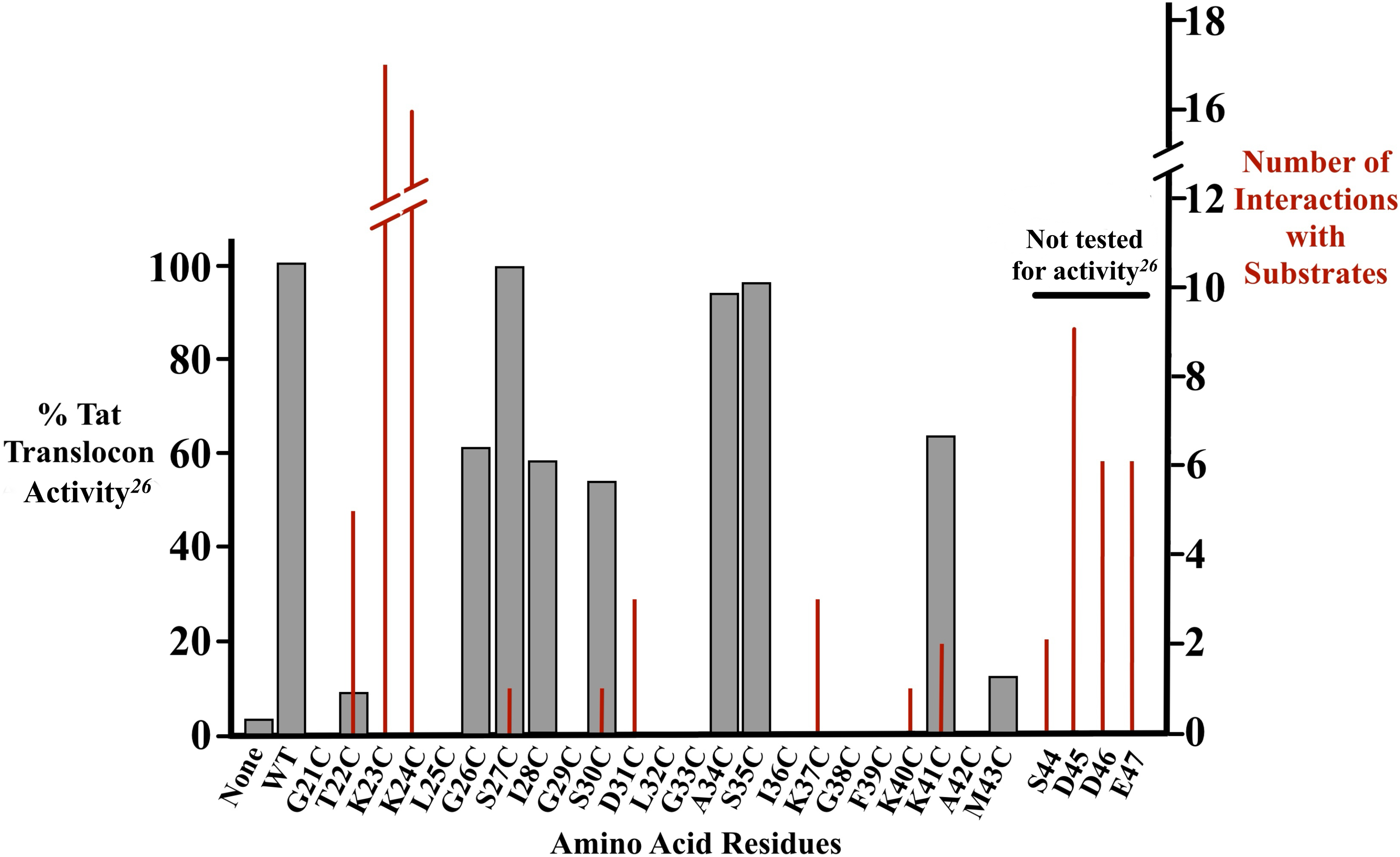
Correlation of the identified substrate-interacting TatA residues with loss of TatA Translocon activity resulting from cysteine-scanning mutagenesis ^26^. Graph of % Tat Translocon activity (left y-axis) vs. amphiphilic helix residues modified by cysteine scanning mutagenesis. This was redrawn from the Greene et al. ^26^, Fig. 2B. The number of apparent charge-pair or apparent hydrogen bond interactions which we observed across both substrates (right y-axis) are shown in red. The TatA residues S44-E47 were not tested for activity in ^26^.

In a higher plant system (*Pisum satvium*), Tha4 (homologue of TatA) residues which are analogous to the TatA region S44-E47 have also been shown to interact with OEE17 ^27^. In this system residues, which had been mutated to cysteine (K46C, E47C, and F48C, corresponding to the TatA residues S44, D45 and D46), were crosslinked to a number of cystinyl-substituted OEE17 mutants (Fig. S3). This result also indicates that TatA residues in this domain directly interact with the OEE17 substrate.

Our simulations in conjunction with the cysteine scanning mutagenesis studies ^26^ and the chemical crosslinking studies ^16, 27^ indicate that substrate-interacting TatA residues were primarily localized in two domains: Domain I (T22-K24) located near the N-terminal portion of the amphipathic helix and Domain II (D45-E47) located at the C-terminal end of the amphipathic helix. Both domains are rich in charged residues (basic and acidic residues, respectively). It should be noted that the clustering of charged residues *per se* is not sufficient to provide sites of interaction between amphipathic helixes and substrates. The region D37-K41 of the amphipathic helix contains multiple charged residues but contributes a relatively small number of amphiphilic helix-substrate interactions.

## Conclusions

The course-grained MD simulations we have described in this communication have proved to be quite useful in examining structure-function relationships of a TatA nonamer and its interaction with Tat Translocon substrates. We believe that our findings provide multiple insights into the mechanistic properties of the role of TatA in the translocation process. First, we demonstrated that the TatA nonamer significantly thins the biological membrane (with or without substrates) (Fig. 2B) confirming earlier MD results ^17^ with the magnitude of the thinning observed being very similar to that previously reported. We also observed numerous phospholipids occupying the lumen pore of the nonamer (Fig 2A). Rodriguez et al. ^17^ described these phospholipids as being “distorted” yielding a presumably structurally altered bilayer. In our simulations these lipids uniformly form a lipid monolayer with the phospholipid head groups facing the p-side of the membrane. While it is unclear if this organization represents the native state of these phospholipids or is an artifact of the MD simulations, the presence of a lipid layer (either a “distorted” bilayer ^17^ or a monolayer) occupying the central pore is necessary to prevent solute leakage across the membrane and would provide a lower energy barrier for substrate translocation. Second, while the TatA nonamer is very unstable in the absence of substrates, in the presence of substrates the oligomer exhibits remarkably enhanced stability. Moreover, the presence of substrate induces a reorganization of the nonamer, leading to the displacement of TatA monomers from the observed radial architecture. This reorganization appears to be related to the size of the substrate, with the larger substrate TorA retaining significantly more TatA monomers than the smaller substrate OEE17. Third, during the time-course of our simulations, both substrates move from the periphery of the TatA nonamer where they are adjacent to the amphipathic helixes of TatA to a more centralized location adjacent to and partially penetrating the oligomer pore while remaining associated with the flexible amphipathic helixes. Simultaneously, water is partially displaced from the n-side of the pore and phospholipids are lost from the oligomer lumen, both of these events would likely be necessary for substrate translocation. We hypothesize that these alterations in TatA oligomer-substrate organization may represent early steps in the translocation process.

Finally, in our simulations the specific hydrophilic residues of the TatA amphipathic helices directly interact with both OEE17 and TorA were identified. These form stable charge-pair and hydrogen-bonding interactions. The amphipathic helix of TatA exhibits robust conformational flexibility particularly at the “hinge” (G21-T22, ^17^) and “elbow” (G32-A33) regions. This conformational flexibility facilitates the direct interaction of hydrophilic amphipathic helix residues with substrates. The interacting TatA residues are clustered primarily in two domains: Domain I (T22-K24) and Domain II (D45-E47), accounting for >75% of the observed interactions. Our results, consequently, complement and extend previous studies using cysteine scanning mutagenesis ^26^ and chemical crosslinking ^16, 39, 59^ to probe the interaction of Tat substrates with TatA.

## Acknowledgements

This work was solely supported by the United States Department of Energy, Office of Basic Energy Sciences grant DE-SC00020304 to SMT and TMB. Special thanks for Ms. Laurie Frankel for providing expert editorial advice.

## Footnotes

1 In this communication we will define “radial architecture” and “radial geometry” as a regular radial distribution of TatA monomers accompanied by a phospholipid-filled pore.

2 In an energized membrane, the n-side of the membrane (in this case the modeled cytoplasm) is more negatively charged than the p-side of the membrane (in this case the modeled periplasmic space). This is due to the pumping of protons to the p-side, with the concomitant formation of !pH and !” which constitute the Proton Motive Force (PMF).

3 For these calculations, if any part of the phospholipid occupied the pore it was included in these calculations. Additionally, CDL1 was counted as 2 phospholipid units.

4 These are termed “apparent” interactions because the individual course-grained beads can represent multiple heavy atoms. This can lead to ambiguity in the direct assignment of the interacting hydrogen- bond and charge-pair atoms.

