## Supplementary Figures and Legends for "Molecular Dynamic Studies on the Interaction of a TatA Oligomer with Tat Translocon Substrates"

Supplementary Figure Legends

Figure S1. Helical wheel of the amphipathic helix of *E. coli* TatA. TatA residue numbers are indicated within the colored circles while numbering within the amphipathic helix are shown outside the circles. The dashed line indicates the boundary between the hydrophobic face (120-140°) and the hydrophilic face (240-220°) of the asymmetric amphipathic helix.

Figure S2. Loss of phospholipids from the oligomer pore during the TatA nonamer + OEE17 and TatA nonamer + TorA simulations. Plot of the number of retained phospholipids vs. trajectory time (ns) for the TatA nonamer + OEE17 (black squares) and TatA nonamer + TorA (red circles) simulations. Each data point is the average observed for 5 simulations.

Figure S3. Sequence alignment of *Pisum satvium* Tha4 and *E. coli* TatA. Color-coding of the domains of TatA and Tha4 are as in Fig. 1. In Tha4, the residues which were observed to be crosslinked to the OEE17 substrate [27] and in TatA, residues which were observed to be crosslinked to TorA and SufI [39] are shown in purple. The residues which we observe to interact with both the OEE17 and TorA, and which are common to these crosslinking studies, are indicated by *.

**Figure S1.**

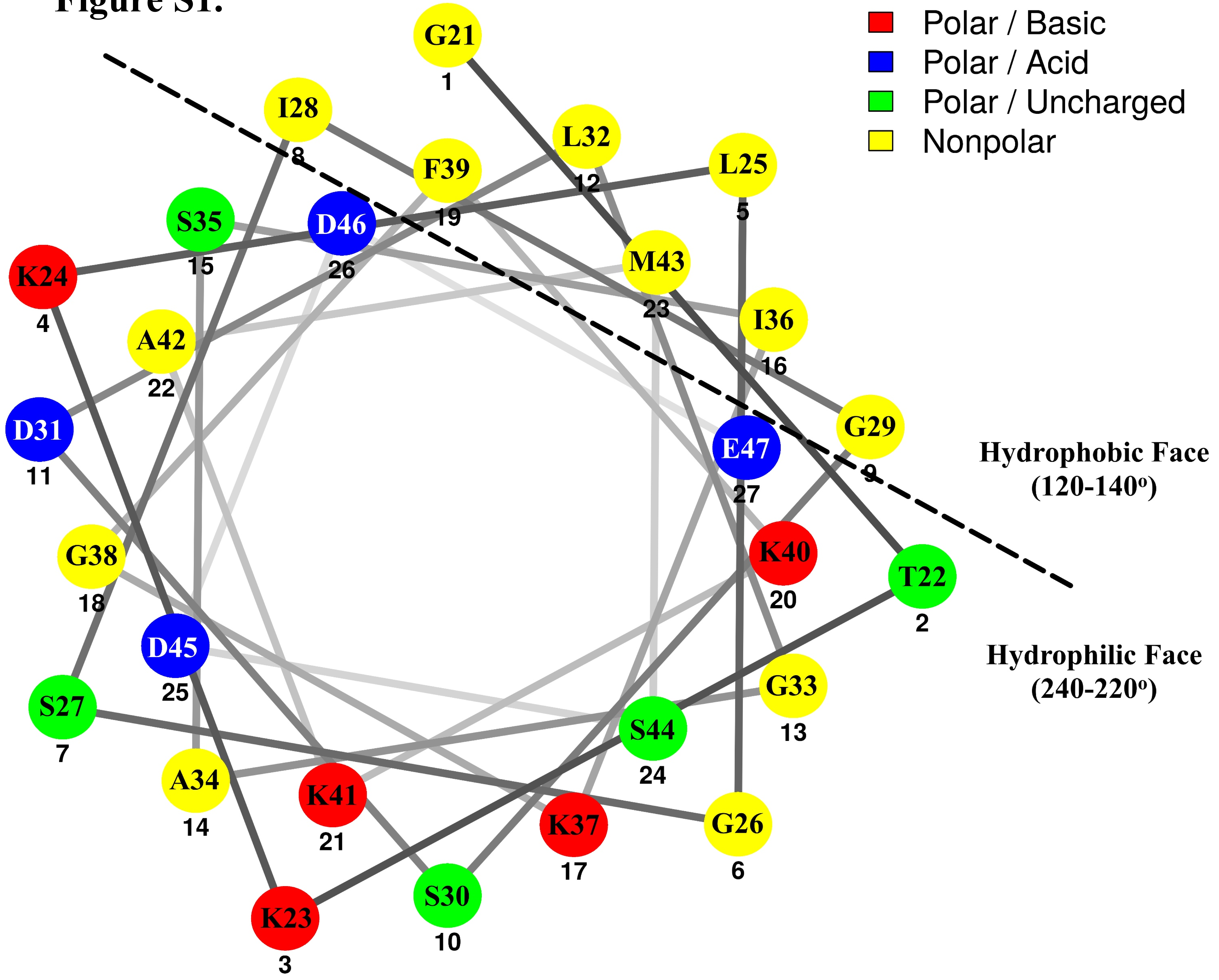

Polar / Basic
Polar / Acid
Polar / Uncharged
Nonpolar
Hydrophobic Face
(120-140°)
Hydrophilic Face
(240-220°)
G21 1
T22 2
K23 3
K24 4
L25 5
G26 6
S27 7
I28 8
G29 9
S30 10
D31 11
L32 12
G33 13
A34 14
S35 15
I36 16
K37 17
G38 18
F39 19
K40 20
K41 21
A42 22
M43 23
S44 24
D45 25
D46 26
E47 27

**Figure S2.**

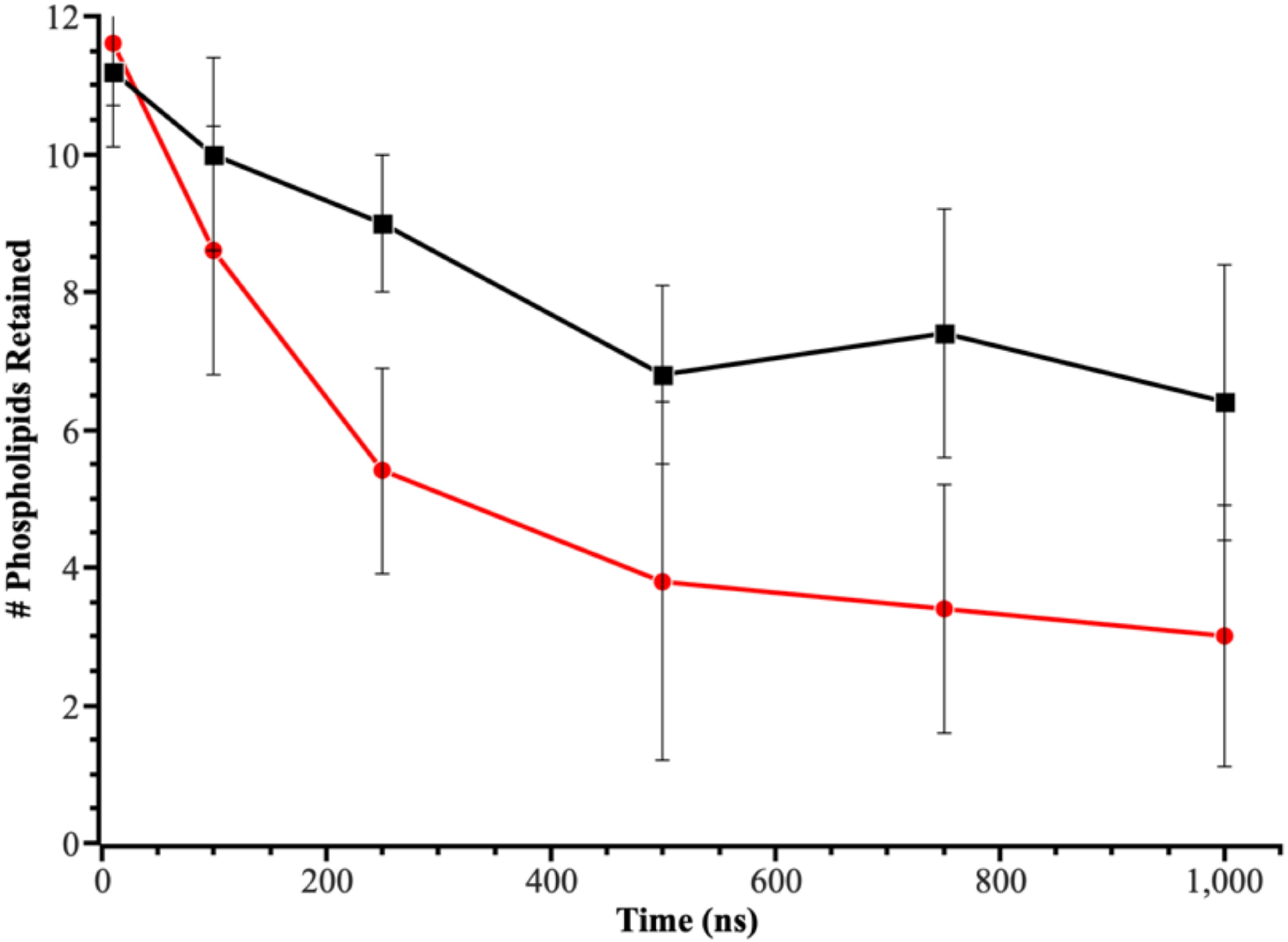

12
10
8
6
4
2
0
### Phospholipids Retained
0
200
400
600
800
1,000
Time (ns)

### Figure S3.

```
                          Transmembrane              Amphipathic
                              Helix                     Helix
                                                *       ***
Tha4      MAFFGLGVPELVVIAGVAALLFGPKKLPEVGRSIGQTVKSFQQAAKEFETELKKEPNPTE 60

TatA      --MGGISIWQLLIIAVIVVLLFGTKKLGSIGSDLGASIKGFKKAMSDDEPKQDKTSQDAD 58
                                      *  **     *        **

Tha4      EI--SVASEQ-------EKQEIKVSSTKDNV 82

TatA      FTAKTIADKQADTNQEQAKTEDAKRHDKEQV 89
              Unstructured C-Terminal
                      Domain
```
